# Snazarus and its human ortholog SNX25 regulate autophagic flux by affecting VAMP8 endocytosis

**DOI:** 10.1101/2021.04.08.439013

**Authors:** Annie Lauzier, Marie-France Bossanyi, Rupali Ugrankar, W. Mike Henne, Steve Jean

## Abstract

Autophagy, the degradation and recycling of cytosolic components in the lysosome, is an essential cellular mechanism. It is a membrane-mediated process that is linked to vesicular trafficking events. The sorting nexin (SNX) protein family controls the sorting of a large array of cargoes, and various SNXs can impact autophagy. To gain a better understanding of their functions *in vivo* under nutrient starvation, we screened all *Drosophila* SNXs by RNAi in the fat body. Significantly, depletion of *snazarus* (*snz*) strongly impacted autolysosome formation and led to decreased autophagic flux. Interestingly, we observed altered distribution of Vamp7-positive vesicles with snz depletion and *snz* roles were conserved in human cells. *SNX25* is the closest ortholog to *snz*, and we demonstrate a role for it in VAMP8 trafficking. We found that this activity was dependent on the *SNX25* PX domain, and independent of *SNX25* anchoring at the ER. We also demonstrate that differentially spliced forms of *SNX14* and *SNX25* are present in cancer cells. This work identifies a conserved role for *snz*/*SNX25* as regulators of autophagic flux, and show differential isoform expression between orthologs.

## Introduction

Macro-autophagy, hereafter termed autophagy, is an essential homeostatic and stress-responsive catabolic mechanism (Doherty and Baehrecke, 2018; Mizushima et al., 2008). Autophagy is characterized by the formation of double membrane structures, called phagophores, which expand and incorporate cytoplasmic proteins or organelles (Gatica et al., 2018; Mizushima et al., 2011). These structures ultimately close to form autophagosomes (Reggiori and Ungermann, 2017). When mature, the autophagosomes fuse with lysosomes, and autophagosomal content is degraded by lysosomal enzymes and recycled (Kriegenburg et al., 2018; Lőrincz and Juhász, 2019; Nakamura and Yoshimori, 2017). Hence, autophagy requires an intricate balance between various cellular processes to ensure appropriate cargo selection, autophagosome formation, maturation, and fusion (Levine and Kroemer, 2019).

While the core signaling pathways controlling autophagy induction in response to stress are well-described (Harder et al., 2014; Korolchuk et al., 2011; Lamb et al., 2013), the molecular mechanisms controlling autophagosome sealing, maturation, and fusion remain unclear. Recent findings in yeast and metazoans have shed light on the molecular machinery required for autophagosome-lysosome fusion and its regulation. Although different proteins are involved in autophagosome-vacuole fusion in yeast and autophagosome-lysosome fusion in metazoans (Reggiori and Ungermann, 2017), the overarching principle is conserved and requires the presence of specific soluble N-ethylmaleimide-sensitive factor attachment receptors (SNAREs) (Itakura et al., 2012; Lőrincz and Juhász, 2019; Nakamura and Yoshimori, 2017). In metazoans, Syntaxin (STX)17 is recruited to mature autophagosomes by two hairpin regions, where it forms a Qabc complex with synaptosome associated protein 29 (SNAP29) (Itakura et al., 2012; Takats et al., 2013). The STX17/SNAP29 complex then forms a fusion-competent complex with lysosome-localized vesicle associated membrane protein (VAMP)8 (Itakura et al., 2012). More recently, the Qa SNARE YKT6 v-SNARE homolog (YKT6) was also found to mediate autophagosome-lysosome fusion (Bas et al., 2018; Matsui et al., 2018; Takáts et al., 2018). YKT6 is recruited to mature autophagosomes and also associates with SNAP29. The YKT6/SNAP29 complex interacts with the lysosomal R-SNARE STX7 to mediate fusion (Bas et al., 2018; Matsui et al., 2018; Takáts et al., 2018). These fusion complexes are conserved, and flies also utilize these proteins for autophagosome-lysosome fusion (Takats et al., 2013; Takáts et al., 2018). However, unlike in human cells, where STX17 and YKT6 act redundantly in parallel pathways, *Ykt6* is epistatic to *Syx17/Vamp7* in flies (Takáts et al., 2018). SNARE functions are supported by other intracellular factors, which ensure their specificity and rapid action (Bröcker et al., 2010; Hong, 2005). The small Rab GTPases Ras-related protein (RAB)7 and RAB2 are important determinants of fusion (Baba et al., 2019; Fujita et al., 2017; Heged s et al., 2016; Kuchitsu et al., 2018; Lörincz et al., 2017; Wang et al., 2016), as lysosomal-localized RAB7 and autophagosomal-localized RAB2 interact with the tethering homotypic fusion and vacuole protein sorting (HOPS) complex to bring autophagosomes and lysosomes in close proximity and enable SNARE-mediated fusion (Fujita et al., 2017; Lörincz et al., 2017; Numrich and Ungermann, 2014). Interestingly, a direct interaction has been observed between STX17 and the HOPS complex (Jiang et al., 2014; Takats et al., 2014), favoring autophagosome-lysosome tethering.

It is clear that multiple inputs are involved and integrated to regulate the final step of the autophagic process. However, VAMP8 and STX7 mediate various other membrane fusion events, and VAMP8 displays minimal localization to endolysosomes in steady-state conditions (Cornick et al., 2019; Diaz-Vera et al., 2017; Jean et al., 2015; Nair-Gupta et al., 2014; Okayama et al., 2009; Pattu et al., 2011; Zhu et al., 2012). This leads to the question of how cells adjust their vesicular routes to account for increased VAMP8 endolysosomal requirement. We previously showed that RAB21 and its guanine nucleotide exchange factor myotubularin-related protein 13 modulate VAMP8 trafficking to endolysosomes upon nutrient starvation (Jean et al., 2015), representing a novel mechanism coordinating autophagy induction with the trafficking of an essential determinant of autophagosome-lysosome fusion. The molecular players involved in this VAMP8 sorting remained to be defined.

One class of endosomal sorting regulators is the sorting nexin (SNX) family (Chi et al., 2015; Cullen, 2008; Van Weering and Cullen, 2014). These proteins have phox homology (PX) domains that interact with diverse phosphoinositide species (Chandra et al., 2019). Many SNXs localize to early endosomes, where they are involved in sorting events (Cullen, 2008). Importantly, a few SNXs play roles in autophagy (Knævelsrud et al., 2013; Maruzs et al., 2015). SNX18 and SNX4 control ATG9 trafficking to modulate autophagosome expansion (Knævelsrud et al., 2013; Ravussin et al., 2021), and SNX5/6 also indirectly regulate autophagy by modulating cation-independent mannose-6-phosphate receptor sorting, affecting lysosomal functions (Cui et al., 2019). However, SNX involvement in the SNARE protein VAMP8 trafficking has not been reported. Here, using *Drosophila* as a simple system to screen genes involved in autophagy, we have identified the sorting nexin *snazarus* (*snz*) and its human ortholog *SNX25* as regulators of Vamp7/VAMP8 localization. Using RNA interference (RNAi) and clustered regularly interspaced short palindromic repeats (CRISPR)/Cas9-generated mutants, we show that loss of *snz* decreases autophagic flux. Importantly, we show that this effect is independent of SNX25’s endoplasmic reticulum (ER) localization and is mediated through its PX domain. Altogether, these findings identify snz and SNX25 as regulators of Vamp7/VAMP8 trafficking, with implications for autophagic flux.

## Results

### *Snz* is required for autophagic flux

To identify SNXs involved in autophagy, we used the *Drosophila* fat body as an inducible system to monitor autophagy *in vivo*. Although SNXs have been previously screened for functions in autophagy, these screens monitored Atg8 and focused on SNXs that decreased autophagosome formation (Knævelsrud et al., 2013; Mauvezin et al., 2016). Thus, to identify SNXs that may have been missed to gene redundancy or weak effects on Atg8, we focused on autolysosomes, using LysoTracker (LyTr) staining of starved third instar fat bodies. LyTr staining has been widely used in the fly fat body (Mauvezin et al., 2014; Rusten et al., 2004; Scott et al., 2004) to assess the terminal steps of autophagy, since autolysosome acidification solely happens under starvation conditions, and in autophagy-proficient cells in this organ (Mauvezin et al., 2014). Using RNAi, we depleted individual SNXs from starved fat bodies and counted the LyTr puncta detected through automated image analysis (Fig. S1A and B). As previously demonstrated, we observed effects for SNX18 (Knævelsrud et al., 2013; Søreng et al., 2018) and most retromer-associated SNXs (Cui et al., 2019), illustrated by significant reductions in the numbers of LyTr puncta in cells depleted of them (Fig. S1A and B). Interestingly, we observed a strong reduction in the number of LyTr puncta in fat bodies depleted of *snz* (Suh et al., 2008; Ugrankar et al., 2019) (Fig. S1A and B). This SNX has four orthologs in humans, *SNX13*, *14*, *19*, and *25* (Henne et al., 2015; Ugrankar et al., 2019). *Snz* and *SNX14* have both been shown to be anchored to the ER, and to mediate lipid droplet (LD) formation in flies and human cells, respectively (Bryant et al., 2018; Datta et al., 2019; Ugrankar et al., 2019). Given the strong autolysosome loss in *snz*-depleted cells and the fact that autophagy and LD formation are intimately linked (Rambold et al., 2015; Velázquez et al., 2016), we examined the role of *snz* in autophagy.

We first analyzed autophagic flux by monitoring ref(2)P, the fly ortholog of p62 (Devorkin and Gorski, 2014). Ref(2)P is an autophagic substrate that is engulfed by autophagosomes and degraded after fusion. Hence, its accumulation suggests defective autophagic flux (Devorkin and Gorski, 2014). Using a validated *snz* RNAi line (Ugrankar et al., 2019) with efficient depletion of ectopically expressed snz (Fig. S1C), we observed increased ref(2)P puncta in snz-depleted fat bodies compared to controls (Fig. 1A and B). This finding was independently corroborated by western blot analysis (Fig. S1D). To more directly assess autophagic flux, we used the green fluorescent protein (GFP):mCherry:Atg8 reporter. This probe allows differentiation between autophagosomes (yellow) and autolysosomes (red), since GFP’s fluorescence is quenched at the acidic pH of the lysosome (Mauvezin et al., 2014). Consistent with the ref(2)P results, we observed an accumulation of autophagosomes (yellow objects) in snz*-*depleted fat bodies compared to control fat bodies (Fig. 1A and C). After CRISPR/Cas9-generated deletion of the *snz* locus (Ugrankar et al., 2019), we observed decreased LyTr puncta (Fig. 1D and E) and autophagosome accumulation (Fig. 1D-F), validating the RNAi results. Given the requirement of snz for full autophagic flux, we tested if snz overexpression could drive or potentiate autophagy. Fat bodies overexpressing an snz:GFP transgene (Ugrankar et al., 2019) did not show an increased abundance of autolysosomes under either fed or starved conditions (Fig. S1E). In contrary, a significant decrease number of LyTr puncta was observed after a three-hour starvation period (Fig. S1E). We conclude from these results that *snz* is necessary but not sufficient for autophagy.

**Figure 1.**
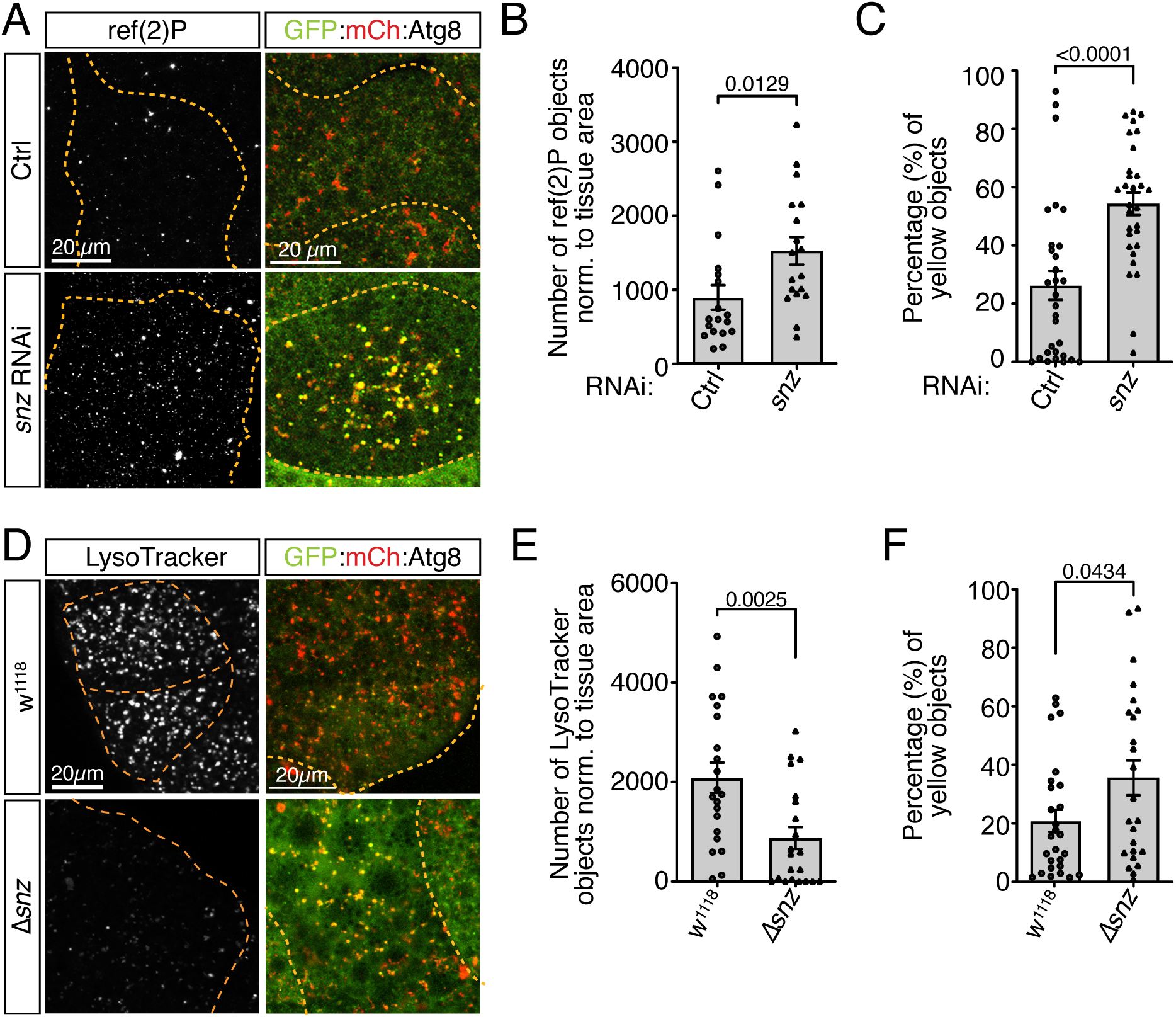
Snz is required for full autophagic flux in the *Drosophila* fat body. Autophagy was assessed in fed and starved (3 h) third instar *Drosophila* fat bodies. (A) Ref(2)P-positive puncta and GFP:mCh:Atg8 colocalization in snz*-*depleted and control fat bodies. (B) Number of ref(2)P- positive puncta normalized to the tissue area. Bars represent the mean, individual points or triangles represent single fat body values, and the error bars are the SEM (*n=*3 independent experiments). (C) Percentage of autophagosomes (yellow objects) per image. Bars represent the mean, individual points or triangles represent single fat body values, and the error bars are the SEM (*n=*4 independent experiments). (D) LyTr-positive puncta and GFP:mCh:Atg8 colocalization in *snz*-mutant and parental (*W^1118^*) fat bodies. (E) Number of LyTr-positive puncta normalized to the tissue area. Bars represent the mean, individual points or triangles represent single fat body values, and the error bars are the SEM (*n=*4 independent experiments). (F) Percentage of autophagosomes (yellow objects) per image. Bars represent the mean, individual points or triangles represent single fat body values, and the error bars are the SEM (*n=*4 independent experiments).

### The autophagic roles of snz are conserved in mammals

*Snz* has four orthologs in mammals, including *SNX14*, a gene mutated in spinocerebellar ataxia autosomal recessive 20 (SCAR20) that can influence autophagy by impacting lysosomal function (Akizu et al., 2015). Sequence alignments indicate that *snz* is more closely related to *SNX25,* and the snz PX domain shows an SNX25-like affinity for diphosphoinositides, rather than the lack of binding specificity displayed by SNX14 (Chandra et al., 2019). To test for functional conservation, we depleted SNX13, SNX14, and SNX25 using two independent small interfering (si)RNAs per gene (Fig. S2A). Analysis of the lipidated form of microtubule associated protein 1 light chain 3 alpha (LC3), LC3-II, revealed increases in SNX14- and SNX25-depleted cells, although these were not statistically significant (Fig. S2B-C). Higher LC3-II levels suggest either increased autophagosome synthesis or defective autophagosome degradation. To assess this, we treated depleted cells with bafilomycin A1 (BafA1) to block autophagosome degradation. In BafA1-treated cells, no increase in LC3-II was observed after SNX-depletion compared to control cells (Fig. S2B). To further strengthen these observations, endogenous LC3 levels were assessed in depleted cells by immunofluorescence. Depletion of SNX14 and SNX25 led to statistically significant increases in the number of LC3 puncta per cell, and no further increase was seen after BafA1 treatment (Fig. 2A and B). These data suggest impaired autophagic flux in siRNA-treated cells. Given the phenotypes observed in SNX14 and SNX25 siRNA-transfected cells, we generated single and double knockout (KO) populations for *SNX14* and *SNX25* using CRISPR/Cas9-based gene editing. Statistically significant increases in LC3-II were observed in independent KO populations (Fig. 2C and D). Altogether, these results argue for positive roles for SNX14 and SNX25 in the control of autophagic flux. They are consistent with the observed effect of snz depletion in the fly fat body and in agreement with findings from patients with SCAR20 (Akizu et al., 2015).

**Figure 2:**
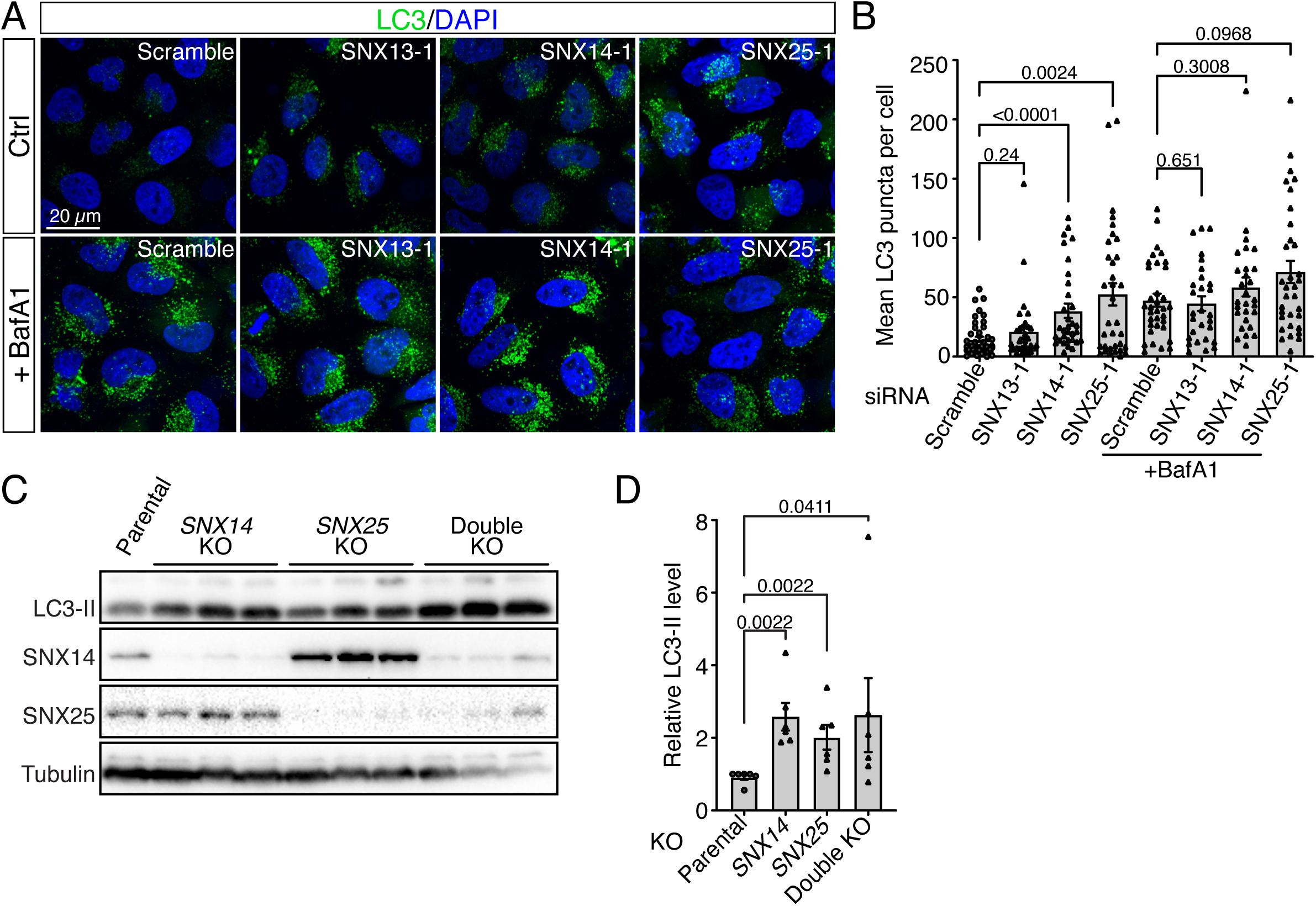
Snz orthologs positively regulate autophagic flux. (A) Immunofluorescence analysis of endogenous LC3 (green) with or without bafilomycin A1 (BafA1) treatment. Nuclei were counterstained with 4′,6-diamidino-2-phenylindole (DAPI). Images are representative of three independent experiments. (B) Quantification of LC3 puncta per cell. Bars represent the mean, individual points or triangles represent single-cell values, and the error bars are the SEM (*n*=3 independent experiments). (C) Western blots of independent *SNX14*, *SNX25,* and double *SNX14* and *SNX25* KO cell populations in normal growth conditions. (D) Quantification of LC3-II levels (normalized to tubulin) in *SNX14*, *SNX25,* and double *SNX14* and *SNX25* KO cell populations. Bars represent the mean of the individual data points and the error bars are the SEM (*n*=6 independent experiments).

### General trafficking pathways are not affected by SNX14, SNX25, or snz depletion

To test which cellular processes were impacted by SNX14 and SNX25 depletion, we first monitored lysosomal functions. Lysosomal cathepsin B activity was assessed using Magic Red, which emits fluorescence only when cleaved by active cathepsin B (Mauvezin et al., 2014). HeLa cells depleted for each *snz* ortholog by siRNA were imaged live using widefield fluorescence microscopy, and the average number of Magic Red puncta was measured in each cell and experiment. A non-significant decrease was observed after SNX14 depletion, while no clear variations were observed in SNX13- or SNX25-depleted cells, suggesting that cathepsin B activity was not impaired (Fig. S2D and E). We also assessed epidermal growth factor receptor (EGFR) degradation in KO cells as an indicator of lysosomal function. As expected, EGF stimulation led to potent extracellular-regulated kinase (ERK) activation and resulted in EGFR degradation in all conditions (Tan et al., 2015), with similar kinetics between control and *SNX14*/*SNX25* single KOs (Fig. S2F and G). Co-ablation of *SNX14* and *SNX25* led to a slight delay in EGFR degradation. Notably, there were no obvious differences in the localization, intensity, and number of early endosome antigen 1 (EEA1)-, adaptor protein, phosphotyrosine interacting with PH domain and leucine zipper 1 (APPL1)-, and CD63 molecule (CD63)-positive compartments (Fig. S3A). From these results, we conclude that *SNX13*, *SNX14,* and *SNX25* can compensate for each other and that their individual loss does not affect lysosomal function; however, co-depletion of *SNX14* and *SNX25* impairs either lysosomal function or EGFR trafficking to the lysosomes to some degree.

Given the partial redundancy observed between *SNX14* and *SNX25* with regard to lysosomal function, we tested the impact of depleting snz, their lone SNX ortholog in flies. RNAi depletion of snz from the fat body did not significantly impact early endosomes, late endosomes, or lysosomes (indicated by Rab5, Rab7, and Lamp1, respectively; Fig. S3B-E). Importantly, both the number and shape of these compartments were similar between control and snz-depleted cells. We also tested lysosomal functions in fly macrophages (hemocytes), given that they have constitutive lysosomal functions, compared to the fat body. LyTr staining and cathepsin B activity did not decrease in the hemocytes of *snz-*ablated flies; on the contrary, increased LyTr- and Magic Red-positive compartments were observed (Fig. S3F-H). Taken together, data obtained from HeLa cells and *Drosophila* indicate that loss of snz or its mammalian orthologs SNX14 and SNX25 impairs autophagic flux quite specifically, without impairing general endo-lysosomal compartments.

### Vamp7/VAMP8 trafficking is perturbed in snz/SNX25-depleted cells

SNARE proteins are essential regulators of autophagic flux (Moreau et al., 2011; Nair et al., 2011) and their trafficking must be regulated to account for increased autophagic demands (Jean et al., 2015). Previous studies have reported trafficking defects for small R-SNAREs that impair autophagy without obvious lysosomal effects (Jean et al., 2015; Moreau et al., 2014). Hence, we assessed the localization of GFP:Vamp7, which represents the single *Drosophila* ortholog of mammalian VAMP7 and VAMP8, in fly fat bodies (Jean et al., 2015; Takats et al., 2013). In control cells, GFP:Vamp7 was observed in plasma membrane (PM) invaginations and in some small intracellular puncta (Fig. 3A, middle panel). In snz-depleted cells, aberrant intracellular GFP:Vamp7 puncta were abundant (Fig. 3A and B, labeled by blue arrowheads in A). Moreover, GFP:Vamp7 aggregates appeared to be in close proximity to the PM in various fat body cells (Fig. 3A, red arrows on the top confocal section and highlighted in the magnified inset). These observations suggest a potential role for snz in modulating Vamp7 internalization or trafficking. To further corroborate these findings, we focused on SNX25, snz’s closest ortholog, and assessed VAMP8 localization in starved *SNX25*-KO HeLa cells. Similar to our observations in flies, VAMP8 accumulated at the PM in *SNX25*-KO cells and displayed reduced colocalization with CD63, a late endosomal protein (Fig. 3C and D). Altogether, these results indicate a conserved role for snz and SNX25 in regulating the trafficking or endocytosis of fly Vamp7 and human VAMP8, respectively.

**Figure 3:**
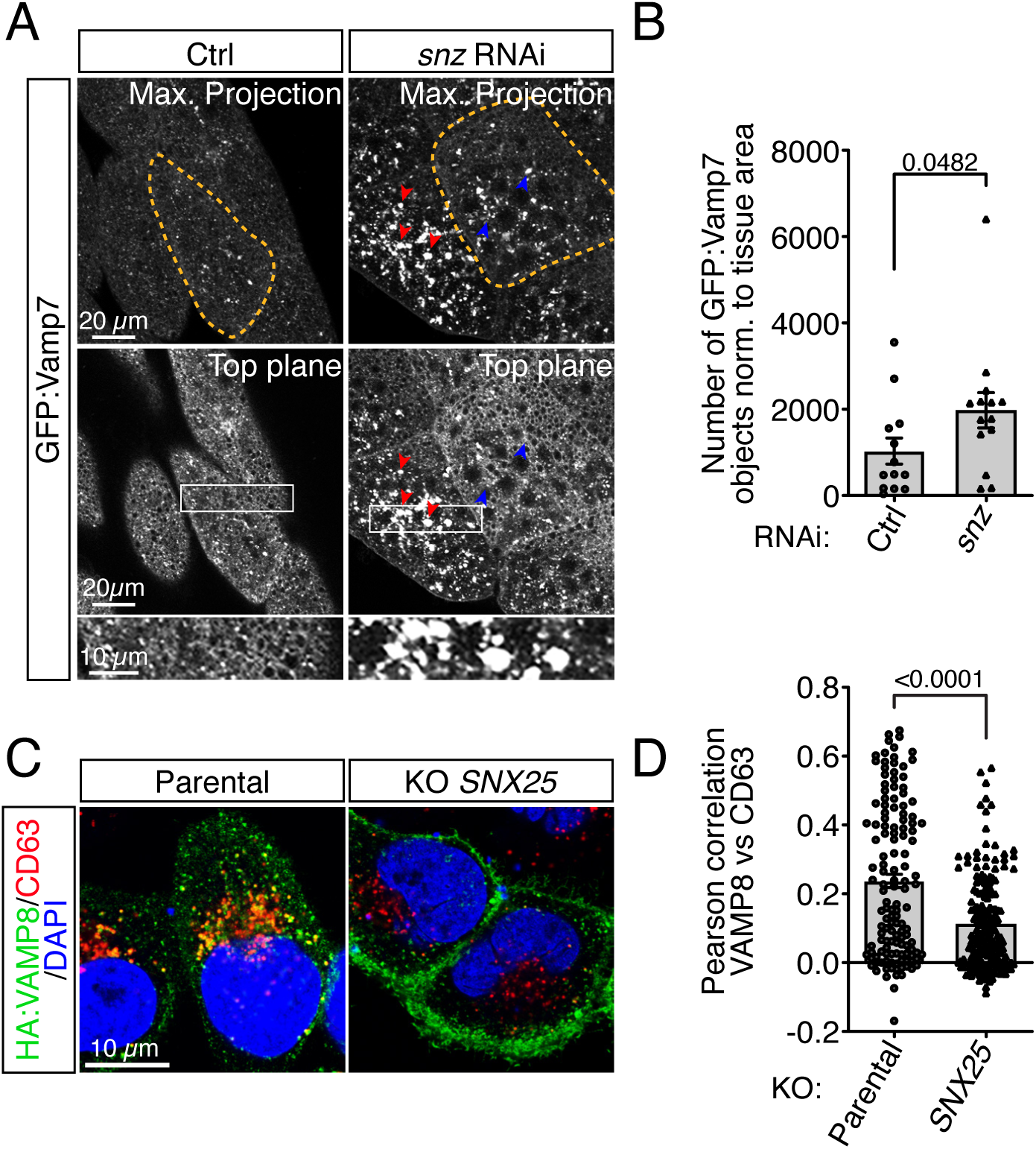
Snz and SNX25 regulate Vamp7/VAMP8 localization. GFP:Vamp7 localization in control and snz-depleted fat bodies. (A) Maximum projections (top panel) and single confocal sections (lower panels) of GFP:Vamp7 puncta in control and snz-depleted cells. Blue arrowheads highlight intracellular puncta, while red arrowheads indicate aggregates present at or near the PM. The bottom panels show magnified views of the boxes in the middle panels. Orange dotted lines indicate single cells. (B) GFP:Vamp7-positive puncta normalized to the tissue area. Bars represent the mean, individual points or triangles represent single data points, and the error bars are the SEM (*n=*3 independent experiments). (C) Parental and KO HeLa cell populations transiently expressing HA-VAMP8 were fixed and stained for endogenous CD63. Nuclei were counterstained with DAPI. (D) Per-cell Pearson correlations between VAMP8 and CD63. Bars represent the mean per-cell correlation, individual points or triangles represent single data points, and the error bars are the SEM (*n*=3 independent experiments).

### Snz and SNX25 colocalize with Vamp7 and VAMP8, respectively

To further examine the link between snz/SNX25 and Vamp7/VAMP8, we performed colocalization studies. It is worth noting that the PX domains of both snz and SNX25 have affinity for diphosphorylated phosphoinositides, including phosphatidylinositol 4,5-bisphosphate (PtdIns(4,5)P2), which localizes to the PM in abundance (Jean and Kiger, 2012b). Previously, snz:GFP was shown to localize close to the PM at ER contact sites, and this localization pattern was dependent on PtdIns(4,5)P2 (Ugrankar et al., 2019). We generated a new snz:mCherry transgenic line, which revealed a punctate localization pattern (Fig. 4A), although it was more diffuse than that produced by snz:GFP (see Fig. S1C). Interestingly, and consistent with the requirement of snz for Vamp7 localization, we observed colocalization between numerous Vamp7- and snz-positive puncta (Fig. 4A and B). Supporting this data, we also observed the highest colocalization, as monitored through Pearson correlation, in HeLa cells between SNX25- HA and GFP-VAMP8 on internal puncta (Fig. S4A and C). Significantly, SNX25 and VAMP8 co-expression in HeLa cells modified their respective localizations, resulting in increased intracellular puncta (Fig. S4A). SNX25 also showed a good degree of colocalization with the ER (indicated by calnexin) and some SNX25 puncta localized to lysosomes and LC3-positive dots. To further assess their proximity, we performed a proximity ligation assay (PLA) between GFP- VAMP8 and SNX25, again in HeLa cells. These experiments confirmed the close proximity between GFP-VAMP8 and SNX25-HA (Fig. 4C and D). As we were unable to detect endogenous SNX25 by immunofluorescence with available antibodies, we performed cellular fractionation experiments to examine the intracellular localization of endogenous SNX25. We observed co-fractionation of SNX25 with VAMP8 and the endosomal marker EEA1 (Fig. 4E). Interestingly, the majority of SNX25 fractionated with endosomal membranes rather than the ER, despite containing two ER-anchoring hairpins. These results highlight the conserved proximity between snz/Vamp7 and SNX25/VAMP8 in flies and humans, and reveal a potential ER-independent pool of SNX25.

**Figure 4:**
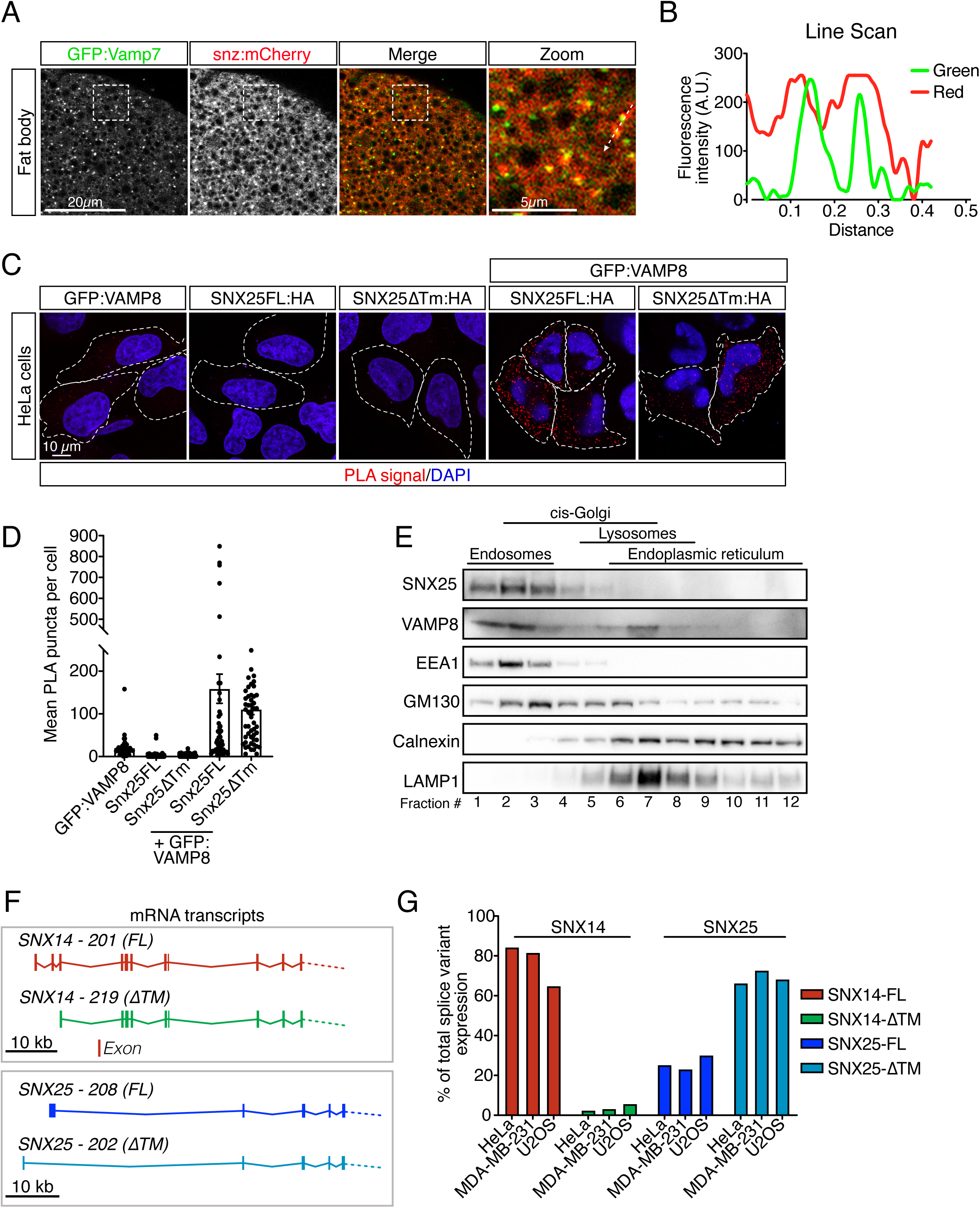
Snz and SNX25 colocalize with Vamp7 and VAMP8 in flies and HeLa cells, respectively. (A) The fat bodies of fed third instar *Drosophila* larvae co-expressing GFP:Vamp7 and snz:mCherry were imaged by confocal microscopy. Images are representative of three independent experiments. A single z-section of the fat body surface is shown. The boxed region has been magnified. (B) Line scan of two GFP:Vamp7 puncta. The path of the line scan is indicated by an arrow in the magnified view in panel A. A.U., arbitrary units. (C) PLA immunofluorescence images showing the proximities between GFP-VAMP8 and SNX25FL-HA or SNX25ΔTM-HA. GFP-VAMP8, SNX25FL-HA, and SNX25ΔTM-HA were also transfected individually and probed with both antibodies as controls (first three images). PLA puncta are shown in red and nuclei were counterstained with DAPI (blue). Dotted lines define individual cells. (D) Quantification of the PLA shown in C. Bars show the mean PLA puncta per cell, individual points represent single cells, and the error bars are the SEM (*n=*3 independent experiments). (E) Immunoblot analysis of SNX25, VAMP8, EEA1, GM130, calnexin, and LAMP1 after a Nycodenz gradient. (F) Schematic representation of the 5′ transcript regions of SNX14 and SNX25. Transcript-associated numbers (from ENSEMBL) are provided, along with the presence or absence of the TM coding sequence. (G) Histogram of the relative transcript levels of FL and ΔTM SNX14/SNX25 isoforms. Multiple smaller transcripts encoding different regions of SNX14 were also quantified but are not depicted in the histogram, which is why the sum of the FL and ΔTM isoforms does not equal 100%.

### Different SNX14 and SNX25 isoforms are expressed in cancer cells

The fractionation result prompted us to assess if *SNX25* (and other family members) had splice variants that could account for the observed discrepancy between the endogenous and exogenously expressed protein localization. In humans, *SNX14* encodes 19 transcripts, with 10 encoding proteins, while *SNX25* has eight transcripts, four of which encode proteins. Strikingly, for both *SNX14* and *SNX25*, specific coding transcripts were identified that encode isoforms lacking ER-anchoring N-terminal transmembrane (TM) domains (Fig. 4F). We thus postulated that the observed differences between our full-length (with ER anchoring) expression construct and the fractionation experiments occurred because the highest abundance SNX25 isoform in HeLa cells may lack the TM domain. Using droplet digital polymerase chain reaction (ddPCR), we quantified the relative abundance of full-length and TM-lacking isoforms in three independent cancer cell lines. In all three cell lines, the SNX25 isoform lacking the TM domain was more abundant than the ER-associated isoform (Fig. 4G). In contrast, SNX14 transcripts mostly encoded longer ER-associated isoforms (Fig. 4G).

Because of the strong preference of the truncated SNX25 isoform lacking the TM domain, we tested its localization. Unlike the ER-anchored full-length SNX25, SNX25-ΔTM was mostly cytoplasmic (Fig. S4B and C). We were not able to observe enrichment at specific sites or a clear colocalization with VAMP8; however, we did observe some proximity by PLA (Fig. 4C and D). From these results, we conclude that in HeLa cells, SNX25 is predominantly not ER-anchored and shows proximity to VAMP8.

### The SNX25 PX domain is necessary for VAMP8 internalization, while ER anchoring is dispensable

The observed accumulation of VAMP8 at the PM in *SNX25* KO cells could be the result of either delayed endocytosis or impaired recycling. To test this more directly, we performed a structure-function analysis of SNX25 in HeLa cells using various truncation mutants (Fig. 5A). Using lentiviral transduction, we expressed SNX25 mutants in *SNX25* KO cells (Fig. S4D) and tracked VAMP8 internalization using an established chase assay (Jean et al., 2015; Miller et al., 2011). It is worth noting that expression of the ΔTM isoform was consistently lower compared to ER-anchored constructs, hinting at a potential faster turnover of cytoplasmic SNX25 (Fig. S4D). *SNX25* KO led to decreased VAMP8 uptake (Fig. 5B and C), suggesting that increased PM VAMP8 is probably due to decreased endocytosis. This was not caused by general endocytic defects, since transferrin internalization (clathrin-dependent) and CD98 uptake (clathrin-independent) were not affected (Fig. S5A-C). Expression of full-length SNX25 rescued VAMP8 uptake, as did the expression of SNX25 mutants missing their TM or C-terminal nexin domains (Fig. 5B and C). This indicates that ER anchoring is not required for SNX25-regulated VAMP8 uptake. However, SNX25 lacking both its PX and nexin domains could not rescue endocytosis (Fig. 5B and C). From the data, we conclude that SNX25 is required for efficient VAMP8 endocytosis, that its anchoring at the ER is dispensable for this regulation, and that its PX domain is required, presumably by mediating SNX25 recruitment to the plasma membrane. These findings are also consistent with the observation that most SNX25 transcripts in various cell types lack the TM domains.

**Figure 5:**
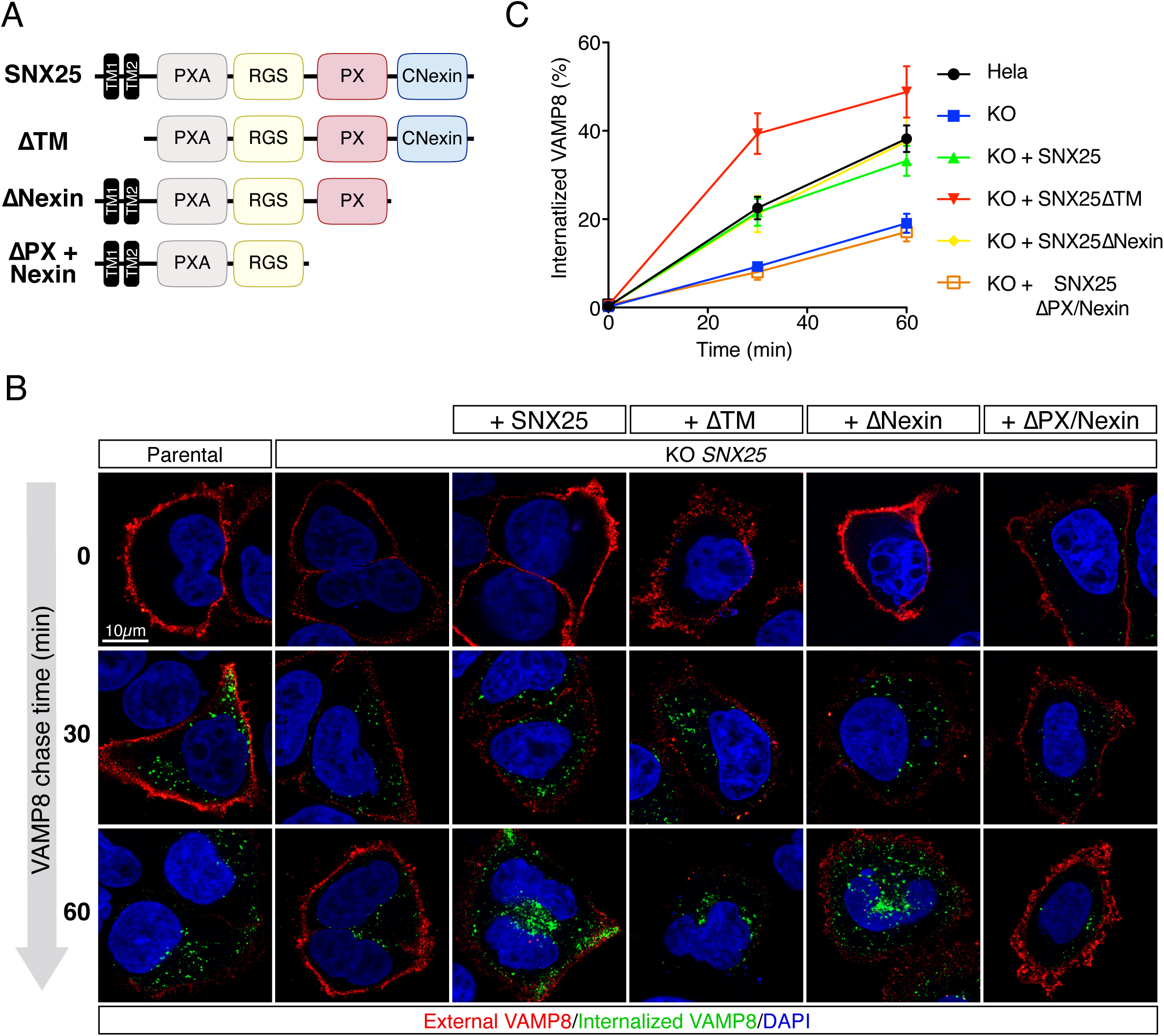
The SNX25 PX domain is required for VAMP8 internalization. (A) Schematic of the SNX25 constructs used for structure/function analysis. (B) VAMP8-GFP uptake in *SNX25* KO cells expressing truncated and full-length forms of SNX25. The graph shows the percentage of the total VAMP8 internalized over time, calculated by measuring the intensity of an external GFP antibody (red) and an internal GFP antibody (green). Nuclei were counterstained with DAPI. Error bars represent the SEM (*n*=3 independent experiments).

### Loss of SNX25 does not affect the levels or recruitment of phosphatidylinositol binding clathrin assembly protein (PICALM) or clathrin

Given the observed effects on VAMP8 uptake in SNX25-deficient cells, we next focused on PICALM, because of its essential role in small R-SNARE internalization (Miller et al., 2011). Moreover, recent proteomic data identified PICALM as a high confidence neighbor of SNX14- APEX2 (Datta et al., 2021). We extended this proteomic finding to SNX25, observing close proximity between SNX25 and PICALM by PLA (Fig. S5D and E). This prompted us to test for the potential misregulation of PICALM in *SNX25* KO cells. Contrary to our hypothesis, no effect on PICALM localization was observed in either *SNX25* KO or *SNX14/SNX25* double KO cells (Fig. S5F and G), ruling out the possibility that defective VAMP8 endocytosis is caused by PICALM dysfunction. As a control of our PICALM membrane staining analysis, we monitored clathrin heavy chain localization and did not observe any reductions in recruitment (Fig. S5F and H), in agreement with the transferrin uptake data. Hence, defective VAMP8 uptake is influenced by SNX25 in a PICALM-independent manner, through a yet-to-be defined mechanism.

## Discussion

Here, we have uncovered a conserved autophagic function for *snz* and its ortholog *SNX25*. Using both RNAi-mediated depletion and CRISPR/Cas9-generated KOs, we show that snz and SNX25 are required for full autophagic flux. We further define that the impact on autophagy is unlikely to occur *via* lysosomal dysfunction, but probably through inappropriate Vamp7 (in flies) and VAMP8 (in humans) internalization and trafficking. Interestingly, the SNX25 PX domain was necessary for VAMP8 uptake, while ER anchoring was dispensable. Moreover, we found that the majority of SNX25 transcripts in three cancer cell lines encoded shorter SNX25 isoforms lacking its TM domains, indicating the potential for functional segregation between LD biogenesis and autophagy regulation.

To further refine endosomal sorting regulators involved in autophagy, we performed a targeted RNAi screen of SNXs in the fly fat body and monitored autolysosome formation (Mauvezin et al., 2014; Rusten et al., 2004; Scott et al., 2004). Unexpectedly, most SNXs tested caused defects in autolysosome acidification. We believe this is a consequence of the wide range of cargos sorted or endocytosed by SNXs (Cullen, 2008). The misrouting of specific cargos could directly or indirectly affect lysosomal function and therefore autolysosome acidification or formation. Our results also reveal the potential for complementation between SNXs paralogs in mammalian cells, which may explain why autophagy defects were not observed for most SNXs in genome-wide screens (Knævelsrud et al., 2013; Moretti et al., 2018; Morita et al., 2018). Autolysosome formation and acidification in the fat body is differentially regulated compared to other tissues. Thus, the observed acidification defect in snz-depleted fat bodies most probably results from the autophagic defects, as seen previously (Jean et al., 2015; Scott et al., 2004), and is independent of endo-lysosomal functions, since multiple assays in either hemocytes or HeLa cells did not detect major defects in endolysosomal compartments and functions.

*SNX14* has three paralogs in mammals, *SNX13*, *SNX19*, and *SNX25*. In neural precursor cells derived from patients with SCAR20, SNX14 loss was associated with autophagosome clearance defects (Akizu et al., 2015). Conversely, weak effects were observed in patients dermal fibroblasts (Bryant et al., 2018). As *Drosophila* have only a single ortholog of these proteins, we were able to show through multiple approaches that loss of snz affected autophagosome clearance and led to autophagosome and autophagic cargo (ref(2)P) accumulation. Our data in HeLa cells also indicate defective autophagic flux in *SNX14*- and *SNX25*-KO cells. The differences between our results and findings in patient fibroblasts might be due to differential regulation of either paralog expression or mRNA splicing between cell types. It is worth mentioning that in HeLa cells, increased SNX14 expression was detected upon *SNX25* KO (Fig. 2C).

Our data indicate defects in the trafficking of Vamp7 and VAMP8 after depletion of snz and SNX25, respectively (Figs. 3 and 4). Since the YKT6/SNAP29/STX7 complex can also promote autophagosome-lysosome fusion, it is likely that this complex partially complements the loss of snz and SNX25, which would explain why their loss did not completely abrogate autophagic flux. Along these lines, differential expression of SNARE complexes between cell types could also account for the variations in penetrance observed between SNX14 studies (Knævelsrud et al., 2013; Moretti et al., 2018; Morita et al., 2018). Finally, under specific stresses, requirements for specific autophagic genes can be bypassed (Kuchitsu et al., 2018). This was recently highlighted for RAB7, when *RAB7* KO cells were shown to be autophagy-proficient under starvation (Kuchitsu et al., 2018). Hence, different starvation protocols or stress sensing capabilities in tested cell types could impact whether SNX14 family members are required for autophagosome clearance.

How exactly snz/SNX25 regulate Vamp7/VAMP8 endocytosis or trafficking remains to be defined. We were unable to directly test Vamp7 trafficking in flies; however, we did observe ectopic accumulation of GFP:Vamp7 puncta near or at the PM, suggesting a potential uptake defect (Fig. 3). To test this more directly, we assessed VAMP8 uptake in *SNX25* KO cells. Interestingly, these cells showed decreased VAMP8 internalization that was dependent on the SNX25 PX domain. This domain interacts with diphosphorylated phosphoinositides (Chandra et al., 2019; Mas et al., 2014) like PtdIns(4,5)P_2_, which is highly abundant at the PM (Jean and Kiger, 2012a). We did not observe defects in clathrin-dependent or -independent endocytosis, nor variations in clathrin recruitment at the PM. Hence, it is unlikely that SNX25 depletion results in VAMP8 trafficking defects by affecting PtdIns(4,5)P_2_ or PtdIns(3,4)P_2_ dynamics at the PM. Recently, snz was demonstrated to bridge PM-ER contact sites to modulate LD formation (Ugrankar et al., 2019). Therefore, SNX25 may fulfill a similar function in mammals, bridging PM-ER contact sites to favor VAMP8 internalization. A precedent for the involvement of ER-PM contact sites in endocytosis exists (Conte et al., 2017); however, we could rescue VAMP8 internalization in *SNX25* KO cells with a transgene lacking its ER-anchoring domains, implying that ER-PM proximity is not required for efficient VAMP8 uptake. This notion is consistent with the known requirement of PICALM for VAMP8 uptake (Miller et al., 2011; Moreau et al., 2014). Surprisingly, we did not detect any defects in PICALM localization in *SNX25* or *SNX14/SNX25* KO cells, although we did observe close proximity between it and overexpressed SNX25. VAMP8 can also be internalized through a clathrin-independent pathway stimulated by Shiga toxin (Renard et al., 2015). This pathway is dependent on lipid organization, and may be perturbed in *SNX25* KO cells. An earlier study identified SNX25 as a regulator of transforming growth factor β receptor (TGFβR) endocytosis. However, this study erroneously characterized the ΔTM isoform of SNX25 and showed that overexpression of this short isoform increased TGFβR internalization (Hao et al., 2011), while SNX25 knockdown decreased uptake. Thus, SNX25 may affect the endocytosis of multiple cargos. This capacity of SNX25 to influence multiple endocytic cargos potentially precluded us from rescuing autophagic defects in SNX25 KO cells through re-expression of the various SNX mutants (data not shown), as done for VAMP8 uptake (Fig. 5). In flies, snz:GFP overexpression partially impaired autophagic flux, and this was also observed in human cells (not shown). Given this, we cannot conclude definitely that the autophagic defect in snz/SNX25 is solely linked to Vamp7/VAMP8 uptake. Nonetheless, our data does argue for a snz/SNX25 role in Vamp7/VAMP8 uptake.

It is also worth mentioning that the yeast ortholog of *snz* and *SNX25*, *MDM1*, was originally identified as a regulator of endocytic trafficking (Henne et al., 2015), thus other aspects of trafficking could be impaired in *snz/SNX25* mutants and be sensitive to protein expression levels. Although our data illustrate decreased internalization of VAMP8 in *SNX25* KO cells, the possibility remains that VAMP8, in addition to its uptake defect, could be misrouted on route to autolysosomes. We did observe decreased colocalization between VAMP8 and CD63 in *SNX25* KO cells; therefore, defective trafficking cannot be ruled out. Moreover, co-expression of both SNX25 and VAMP8 led to the re-localization of both proteins to large internal vesicles. This effect required the TM region of SNX25, thus it is conceivable that although the short isoform is sufficient for VAMP8 internalization, the longer ER-associated isoform could regulate the endosomal sorting of VAMP8, through potential interorganelle contact sites.

Recent studies have demonstrated important roles for SNX14 in lipid metabolism (Bryant et al., 2018; Datta et al., 2019; Datta et al., 2021; Hariri et al., 2019). SNX14 loss results in saturated fatty acid accumulation and increased sensitivity to lipotoxic stress. Moreover, SNX14, snz, and the yeast ortholog were all shown to regulate LD formation (Bryant et al., 2018; Datta et al., 2019; Hariri et al., 2018; Hariri et al., 2019; Ugrankar et al., 2019). Interestingly, the functional domains required for SNX14 regulation of LD formation differ from the ones required in SNX25 for VAMP8 uptake: the TM and C-terminal nexin domains of SNX14 are essential for LD localization and regulation (Datta et al., 2019), while the PX domain of SNX25 is required for VAMP8 uptake, and its TM domains are dispensable. Interestingly, LD biogenesis, FA trafficking, and autophagy are known to intersect (Nguyen et al., 2017; Rambold et al., 2015). In this context, it is tempting to speculate that snz and its human orthologs SNX14 and SNX25 could bridge lipid stress and autophagy regulation. How exactly these proteins favor one process over the other will be an interesting question to address.

The observation that various isoforms of SNX14 and SNX25 are expressed in cells is intriguing. This highlights the possibility of functional pools of SNX14 and SNX25, with the longer ER-anchored isoform regulating LD biogenesis and the shorter isoforms regulating other processes, like trafficking and autophagy. It is worth noting, however, that although we provide evidence from ddPCR experiments, we were unable to demonstrate differential splicing at the protein level because of a lack of isoform-specific antibodies. Isoform expression may be controlled by modulating splicing in response to stress, as has been observed for multiple genes (Biamonti and Caceres, 2009). Alternatively, different transcription factors may favor the expression of certain isoforms (Babeu et al., 2018). RNA-sequencing datasets from *Drosophila* do not contain different snz isoforms, suggesting that a single isoform regulates both LD biogenesis and autophagy.

In summary, we have identified a new role for snz and its ortholog SNX25 in autophagy regulation through Vamp7/VAMP8 internalization and described differentially expressed isoforms of SNX14 and SNX25 in cancer cells. Based on our results and those of previous studies (Akizu et al., 2015; Bryant et al., 2018), we propose that snz and SNX25 fine-tune the endocytosis/trafficking of Vamp7 and VAMP8 to coordinate the level of autophagy with the cell’s demands, and potentially balance the process with lipid handling. It will be interesting to define how these functions are adjusted between various genes and isoforms, and how they are affected by different stressors.

### Experimental Procedures

#### Drosophila strains

Cg-Gal4 was used to drive UASt-RNAi hairpin expression in the fat body and hemocytes. Genotypes used in this study include: (1) *w; UAS-IR-Snz^V105671^*, (2) *Δsnazarus* (From (Ugrankar et al., 2019)), (3) *w; UASp-GFP:mCherry:Atg8a* (from I. Nezis and H. Stenmark), (4) *y^1^ sc* v^1^ sev^21^; P{TRiP.GL01875}attP40* (Bloomington 67953; SNX1 RNAi, (5) *w; UAS-IR- SNX3^V104494^*, (6) *w; UAS-IR-SNX6^V24275^*, (7) *w; UAS-IR-SNX17^V109452^*, (8) *w; UAS-IR-SNX18^V22412^*, (9) *w; UAS-IR-SNX21^V101320^*, (10) *w; UAS-IR-SNX23^V40603^*, (11) *w; UAS-IR-SNX27^V108542^*, (12) *w; UAS-Snz:GFP* (from (Ugrankar et al., 2019)), (13) *w; Cg-GAL4, UAS-GFP:Vamp7^3^/ CyO* (From (Jean et al., 2015)), (14) *w; Cg-GAL4, UAS-GFP:Lamp1* (From (Jean et al., 2015)), (15) *w; Cg- GAL4, UAS-GFP:Rab7^2^* (From (Jean et al., 2015)), and (16) *w; Cg-GAL4, UAS-GFP:Rab5* (From (Jean et al., 2015)).

New lines generated in this study were: (1) *w; Cg-Gal4; p{w[mC] = UASt-Snz:GFP}*, (2) *w; p{w[mC] = UASt-Snz:mCherry}*.

### Drosophila crosses and starvation protocol

Fly stocks were maintained on standard cornmeal food (Bloomington). Fly crosses were housed at room temperature for 2 d, then shifted to 29°C for 3 d. To ensure that third instar larvae were not crowded, 20-30 larvae were transferred to new vials 16-20 h before experiments. Actively feeding third instar larvae were starved by incubating them on Kimwipes prewetted with 1× phosphate-buffered saline (PBS) for 3 h at 25°C.

### Immunofluorescence, uptake assays, PLA, and microscopy

Fat bodies were imaged live and dissected in 1× PBS. For LyTr experiments, fat bodies were incubated in a 1:5,000 dilution in 1× PBS of LyTr for 5 min at room temperature and washed once in 1× PBS. For all other conditions, fat bodies were dissected in 1× PBS only. Fat bodies were mounted on #1.5 coverslips using silicone grease and five individual fat bodies per condition were imaged per experiment. For ref(2)P experiments, larvae were inverted and fixed for at least 2 h in 1× PBS containing 8% paraformaldehyde. Larvae were blocked and permeabilized by a 2 h incubation at room temperature in 1× PBS containing 5% goat serum, 1% bovine serum albumin (BSA), and 0.3% Triton X-100. Larvae were incubated overnight at 4°C in 1:500 anti-ref(2)P antibody (Abcam, ab178440) diluted in 1× PBS containing 1% BSA and 0.3% Triton X-100. Larvae were washed five times for 5 min each in 1× PBS at room temperature and incubated with secondary antibodies (1:250, anti-rabbit Alexa Fluor 488, Life Technologies) for 2 h at room temperature. Larvae were washed as above, and fat bodies were dissected and mounted in 1× PBS for imaging.

Wild type, siRNA-treated, and KO HeLa cells were seeded on #1.5 coverslips and treated according to the various experimental schemes. Cells were fixed in 4% paraformaldehyde and immunofluorescence was performed following the Cell Signaling Technology protocol. Magic Red (ImmunoChemistry Technologies, Bloomington, MN, USA) staining was performed following the manufacturer’s instructions, and live cells were analyzed on a Celldiscoverer 7 (Zeiss) equipped with an environmental chamber. Antibodies used for immunofluorescence were: anti-GFP (1:500, MilliporeSigma, 11814460001), anti-HA (1:1,000, Cell Signaling Technology, Danvers, MA, USA, 3724), anti-CD63 (1:250, BD Biosciences, 561983), anti-LC3 (1:1000, Cell Signaling Technology, 12741), anti-APPL1 (1:200, Cell Signaling Technology, 3858), anti-EEA1 (1:100, Cell Signaling Technology, 3288), anti-calnexin (1:50, Cell Signaling Technology, 2679), anti-GM130 (1:3,000, Cell Signaling Technology, 12480), anti-PICALM (1:200, Abcam ab172962), and anti-clathrin heavy chain (1:50, Cell Signaling Technology, 4796). Secondary antibodies (anti-mouse Alexa Fluor 546 (#A11003), anti-rabbit Alexa Fluor 546 (#A11035), anti-mouse Alexa Fluor 488 (#A11029), and anti-rabbit Alexa Fluor 488 (#A11008) were purchased from Thermo Fisher Scientific (Waltham, MA, USA) and used at 1:500. Cells were counterstained with DAPI (1:10,000, Cell Signaling Technology)

PLA experiments were performed as previously described (Del Olmo et al., 2019b), except that the primary antibodies used for the PLA reaction were anti-HA (1:1,000, Santa Cruz Biotechnology, Dallas, TX, USA, 7392) and anti-GFP (1:500, Thermo Fisher Scientific, A6455). Transferrin and CD98 uptake assays were performed as previously described (Del Olmo et al., 2019a). Finally, VAMP8 uptake assays were performed as described (Jean and Kiger, 2016) except that VAMP8-GFP was used instead of VAMP8-3×HA. A 1:100 dilution of anti-GFP (MilliporeSigma, 11814460001) was diluted in Dulbecco’s modified Eagle’s medium (DMEM) supplemented with 10% fetal bovine serum (Wisent, Saint-Bruno, QC, Canada) and 1% penicillin/streptomycin (complete DMEM) and used to chase VAMP8-GFP uptake. Secondary antibodies used for uptake experiments were anti-mouse Alexa Fluor 555 (#A32727) and anti-mouse Alexa Fluor 647, (#A32728) from ThermoFisher.

Images were acquired on an Olympus FV1000 microscope with a UPlanSApo 40× 1.3 NA oil objective (in Fig. 1A and D, Fig. 3A, Fig. S1A and E, Fig. S1E, and Fig. S3B), a Zeiss LSM 880 microscope with a W PlanAPO 40× 1.4 NA oil objective (Fig. 2A, Fig. 3C, Fig. 4A and C, Fig. 5B, Fig. S1C, Fig. S3A and F, Fig S4A and B, and Fig. S5A, D, and F), or a Celldiscoverer 7 using a Plan-Apochromat 20× WD 0.8 NA objective (Fig. S2D). Control samples were imaged first to set acquisition parameters, which were conserved for all samples in each experimental set. Images were exported to .tiff files and quantified using Cell Profiler as previously described (Jean et al., 2015). Images were prepared for publication using Photoshop 2021 (Adobe, San Jose, CA, USA). Only linear level adjustments were performed, which were kept consistent between experimental sets. Enhanced images were cropped and merged in Photoshop and assembled in Adobe Illustrator.

### Generation of DNA constructs

Full-length snz:GFP was used to generate the snz:mCherry construct. Briefly, the *snz* coding region (from snz:GFP) and mCherry (from pmCherry-C1) were PCR amplified and ligated into the pUASt-Attb plasmid using the In-FusionHD Cloning Kit (Takara Bio, Mountain View, CA, USA). The plasmid was injected into flies. Full-length *SNX25* was PCR amplified from HeLa cell cDNA generated with the Superscript III First-Strand Synthesis System (Thermo Fisher Scientific) and cloned into pcDNA3-3×HA using the In-FusionHD Cloning Kit. This plasmid was used as a template to generate the various truncation mutants. Lentiviral vectors were generated by subcloning the different pcDNA3-SNX25-HA mutants into a BamHI/AfeI-digested pLVX vector using the In-FusionHD Cloning Kit. All vectors generated were validated by sequencing.

### Cell culture, CRISPR/Cas9 KO, and lentiviral rescue

HeLa cells were maintained in complete DMEM at 37°C and 5% CO2. Transient plasmid and siRNA transfections were performed with JetPrime (PolyPlus-transfection, New York, NY, USA) and Dharmafect (Dharmacon, Lafayette, CO, USA), respectively, according to the manufacturers’ instructions. Cells were analyzed by western blot or immunofluorescence 24 h after plasmid transfection and 72 h after siRNA transfection. The siRNAs used were SNX13-1 (J- 009381-10-0002), SNX13-2 (J-009381-11-0002), SNX14-1 (J-013190-10-0002), SNX14-2 (J-013190-12-0002), SNX25-1 (J-014761-11-0002), SNX25-2 (J-014761-12-0002), and a non-targeting control (scramble, D-001810-01-05), at a final concentration of 10 nM. Knockdown efficiencies were measured by quantitative reverse-transcription (qRT)-PCR, and relative mRNA levels were calculated using the ΔΔCT method (Livak and Schmittgen, 2001) and normalized to glyceraldehyde 3-phosphate dehydrogenase (GAPDH) (Del Olmo et al., 2019b).

KO HeLa cell populations were generated as previously described (Del Olmo et al., 2019a). Briefly, three independent gRNAs per gene (sequences obtained from (Sullender et al., 2016)) were cloned into pX330A and validated by sequencing. Cells were transfected with the three gRNAs along with the pEGFP-Puro plasmid using JetPrime at a 10:1 ratio (300 ng of each pX330A-gRNA and 100 ng of pEGFP-Puro). The following day, transfected cells were selected for 36 h with puromycin (1µg/ml), followed by expansion in complete DMEM for two passages. KO efficiencies were then validated by western blot analysis. KO cell populations were used at low passage numbers for all experiments, to ensure that residual wild-type cells would not outcompete edited cells.

For KO/rescue experiments, lentiviruses were produced in 293T cells and supernatants were collected and stored. *SNX25* KO cells were transduced with the various lentiviruses and selected with puromycin (1µg/ml) for 2 d. Following selection, cells were expanded for one passage and used in experiments. Expression of the various rescue constructs was confirmed by western blot analysis (Fig. S4D).

### Western blot analysis

To analyze autophagy, HeLa cells (5×10^4^ cells for siRNA transfections or 2×10^5^ KO cells) were seeded in individual wells of a 24-well plate. Cells transfected with siRNA were processed 72 h after transfection, while KO cells were processed 24 h after plating. Cells were lysed in 150 µL radioimmunoprecipitation (RIPA) buffer (20 mM Tris-HCl pH 7.5, 150 mM NaCl, 1 mM ethylenediaminetetraacetic acid (EDTA), 1 mM ethylene glycol tetraacetic acid, 1% NP-40, 0.1% sodium dodecyl sulfate (SDS), and 2× protease inhibitors (SigmaAldrich)) for 20 minutes on ice. Protein extracts were centrifuged at 13,200 rpm for 10 min at 4°C. Supernatants were collected and quantified using the Pierce BCA Protein Assay Kit (Thermo Fisher Scientific). Equal amounts of proteins were resolved by SDS-polyacrylamide gel electrophoresis (PAGE) and transferred to PVDF (Immobilon) membranes. The following antibodies were used for western blot analysis: anti-LC3 (1:1,000, Cell Signaling Technology, 12741), anti-SNX14 (1:500, Sigma-Aldrich, St. Louis, MO, USA), hpa017639), anti-SNX25 (1:500, Abcam, ab183756), anti-tubulin (1:2,500, Sigma-Aldrich, T9026), anti-VAMP8 (1:1,000, Synaptic Systems, Göttingen, Germany, 104302), anti-EEA1 (1:1,000, Cell Signaling Technology, 3288), anti-GM130 (1:1,000, Cell Signaling Technology, 12480), anti-calnexin (1:1,000, Cell Signaling Technology, 2679), anti-LAMP1 (1:1,000, Cell Signaling Technology, 9091), anti-ref(2)P (1:500, Abcam, ab178440), anti-GAPDH (1:2000, Cell Signaling Technology, 8884), anti-EGFR (1:1,000, Cell Signaling Technology, 4267), anti-phosphorylated-ERK Thr 202/204 (1:1,000, Cell Signaling Technology, 4370), and anti-HA (1:1,000, Cell Signaling Technology, 3724). Horseradish peroxidase-coupled secondary antibodies were purchased from Jackson ImmunoResearch (West Grove, PA, USA). Bands were detected by chemiluminescence using Luminata Forte (Millipore).

Analysis of ref(2)P (a kind gift from G. Juhasz) in fat bodies was performed as described (Jean et al., 2015) with a few modifications. Briefly, five third instar control or snz RNAi-expressing larvae were dissected, and all fat body tissues removed. Fat bodies were lysed directly in 100 µL of 1× Laemmli buffer for 20 minutes at 4°C. Fat body extracts were centrifuged at 13,200 rpm for 10 min at 4°C. Each supernatant was collected and 25 µL was loaded directly on an 8% SDS-PAGE gel and analyzed by western blotting as described above.

### Protein fractionation

Cell fractionation was performed as previously described (Reekmans et al., 2010) with some modifications. Briefly, for each sample, two confluent 150 mm plates of HeLa cells were washed twice with ice-cold 1× PBS, trypsinized, and resuspended in DMEM. Cells were centrifuged at 500 × *g* for 5 min and washed twice in ice-cold 1× PBS. Each cell pellet was resuspended in 10 mM triethanolamine (pH 7.4), 10 mM acetic acid, 250 mM sucrose, 1 mM

EDTA, 1 mM dithiothreitol, and 1× protease inhibitor. Cells were homogenized using a Dounce homogenizer and centrifuged at 500 × *g* for 10 min at 4°C. The supernatant was collected and loaded on a 10–25% Nycodenz (SigmaAldrich) continuous gradient. The gradient was centrifuged for 90 min at 170,000 × *<u>g</u>* in a swinging bucket rotor. Twelve 1-mL fractions were collected from the top to the bottom of the gradient using a P1000 manual pipettor. Fractions were precipitated using trichloroacetic acid and resuspended in 100 µL of 1× Laemmli buffer, and 25 µL was resolved by SDS-PAGE and analyzed as described above.

### EGFR degradation

HeLa cell KO populations were cultured to 80% confluence, serum-starved for 16 h, and treated with 100 ng/mL EGF and 25 µg/mL cycloheximide for the indicated period of time (Jean et al., 2015). Cells were then lysed in RIPA buffer, processed, and analyzed as described above.

### ddPCR

Primers were designed as described elsewhere (Brosseau et al., 2010). All primers were individually resuspended to 20–100μM in Tris-EDTA buffer (IDT) and diluted as primer pairs to 1 μM in RNase DNase-free water (IDT). Primer validation was performed on a CFX96 Real-Time PCR Detection System (Bio-Rad) with 5 μL of 2× PerfeCTa SYBR Green SuperMix Reagent (Quantabio, Beverly, MA, USA). Amplified products were analyzed by automated chip-based microcapillary electrophoresis on a Labchip GX Touch HT Nucleic Acid Analyzer (Perkin Elmer). Primer sequences are listed in Supplementary Table 1.

For ddPCR, reactions were composed of 10 μL of 2× QX200 ddPCR EvaGreen Supermix (Bio-Rad), 3 µL (150 ng) cDNA, 4 µL paired primers (final concentration: 200 nM), and 3 µL molecular grade sterile water (Wisent) in a 20 µL total reaction. Reactions were converted to droplets with the QX200 Droplet Generator (Bio-Rad). Droplet-partitioned samples were then transferred to a 96-well plate. The plate was sealed, and ddPCR was performed in a C1000 Touch Thermal Cycler (Bio-Rad). The plate was then transferred to a QX200 Droplet Reader and read (Bio-Rad). The concentrations (in copies/µL) of the final 1× ddPCR reactions were determined using QuantaSoft software (Bio-Rad) and the abundance of each transcript was normalized to total transcripts abundances.

### Experimental design and statistical analysis

The RNAi screen presented in Fig. S1A was performed in independent duplicates for each RNAi reaction. Five genes were screened along with a starved control. Independent data sets were pooled and graphed as in Fig. S1B. All other experiments performed in the fly fat body were performed in at least independent triplicates. All experiments performed in HeLa cells also involved at least three independent replicates.

Statistical analyses were performed using GraphPad Prism (GraphPad Software, San Diego, CA, USA) and all graphs show individual data points and the standard error of the mean (SEM) to ease the analysis of experimental variations. For some quantifications, where curves are depicted, only SEMs are displayed. For statistical analyses, normality was first assessed using a D’Agostino-Pearson omnibus normality test. Samples with normal distributions were analyzed by unpaired *t*-tests, while nonparametric Mann-Whitney tests were performed for samples with non-normal distributions.

## Acknowledgments

We thank members of the Jean laboratory for their helpful opinions during this work and Amy Kiger for her support in establishing tools for this study. We thank Alexander Sorkin for generously providing the GFP:PICALM plasmid. We thank the Photonic microscopy platform for confocal use, and the RNomics platform for primer design and for performing ddPCR reactions. We thank High-Fidelity Science Communications for editing the manuscript. Steve Jean is a member of the Fonds de Recherche du Québec - Santé (FRQS)-Funded Centre de Recherche du CHUS and is a recipient of a Research Chair from the Centre de recherche médicale de l’Université de Sherbrooke (CRMUS). This research was supported by an operating grant from the Canadian Institutes of Health Research (CIHR). Marie-France Bossanyi was supported by master’s degree fellowships from the Natural Sciences and Engineering Research Council of Canada and FRQS. Steve Jean is supported by junior faculty salary awards from CIHR and FQRS (FRQS-J1 and J2).

## Author contributions

AL, MFB, and SJ designed, analyzed, and performed the experiments. RU and MWH provided reagents and suggestions throughout this work. SJ wrote the manuscript.

